# Cortical Tonic Inhibition Regulates the Expression of Spike-and-Wave Discharges Associated with Absence Epilepsy

**DOI:** 10.1101/164947

**Authors:** Kile P. Mangan, Aaron B. Nelson, Steven Petrou, Chiara Cirelli, Mathew V. Jones

## Abstract

Synchronous and bilateral spike-and-wave discharges accompany nonconvulsive behavioral and cognitive arrest during seizures associated with absence epilepsy. Previous investigation of multiple absence animal models suggests that the underlying cause of absence seizures is an increase in thalamic inhibitory tonic currents. In contrast, in this study we provide evidence that the level of cortical tonic inhibition also regulates absence seizure expression. Using continuous video-EEG recordings to monitor absence seizures and spike-and-wave discharge expression we show that pharmacological blockade of cortical tonic inhibition provokes absence seizures in wild-type mice. Furthermore, we show that pharmacological rescue of cortical tonic inhibition in an absence mouse (γ2R43Q) model, which lacks tonic inhibition, suppresses absence seizure and spike-and-wave discharge expression. Collectively, these results suggest an optimum level of tonic inhibition in the thalamocortical circuit is required for normal functioning and that a deviation from this optimum results in aberrant thalamocortical function, SWDs and absence seizures.

## INTRODUCTION

Absence seizures are characterized by periods of behavioral arrest, accompanied by glassy staring with loss of cognitive function, often without overt convulsions (McCormick and Contreras, 2001). These generalized seizures include a brief loss of consciousness typically lasting 2-15 s with corresponding bilateral, synchronous ∼3 Hz electrographic spike-and-wave discharges (SWDs) (Panayiotopoulos et al., 2008). The generalized nature of absences, however, should not be rigidly interpreted to mean that there is no specific focus or generating region. SWD initiation has been localized to the perioral region of the somatosensory cortex in multiple animal models of absence epilepsy (Meeren et al., 2002; Meeren et al., 2005; van Luijtelaar & Sitnikova, 2006; van Luijtelaar et al., 2006; van Luijtelaar et al., 2011; L’uttjohann & van Luijtelaar, 2012; Pinault, 2003; Polack et al., 2007; Bazyan & van Luijtelaar, 2013) and local injection of different pharmaceutical agents into this area suppresses absence seizures in multiple SWD-expressing animal models (Manning et al., 2004; Chen et al., 2011; Bazyan & van Luijtelaar, 2013). A “cortical focus theory” for absence seizures has been proposed and many absence animal models support this conclusion (Meeren et al., 2005; Steriade and Contreras, 1998; Sitnikova and van Luijtelaar, 2004; van Luijtelaar and Sitnikova, 2006; Polack et al., 2007).

The γ2R43Q mutation, an arginine-to-glutamine substitution at position 43 of the GABA_A_ receptor γ2 subunit, confers absence epilepsy and febrile seizures in humans (Wallace et al., 2001). Compared to unaffected family members, patients harboring the γ2R43Q mutation display increased intracortical excitability, decreased intracortical inhibition, and increased cortical facilitation (Fedi et al., 2008). Similarly, γ2R43Q knock-in (RQ) mice display absence seizures, generalized SWDs (∼6 Hz), and increased cortical excitability (Tan et al., 2007; Mangan et al., 2013). Results from several labs conclude that the γ2R43Q mutation alters the membrane trafficking of several GABA_A_ receptor subunits to the cell surface (Kang & Macdonald, 2004; Sancar & Czajkowski, 2004; Hales et al., 2005; Eugene et al., 2007; Mangan et al., 2013), including the α5 (Eugene et al., 2007) and δ (Mangan et al., 2013) subunits in cortex and thalamus, respectively. The altered trafficking of these specific subunits results in the loss of GABAergic tonic currents in principal neurons of the thalamocortical loop, leading to increased cortical excitability and decreased thalamic bursting behaviors (Mangan et al., 2013).

Recently a link has been established between tonic inhibition and absence-associated SWD generation, though this link is currently isolated to, and believed to be ‘required’ in, thalamic relay neurons (Fariello & Golden, 1987; Crunelli et al., 2011; Errington et al., 2011). Evidence of this link includes findings that the *increase* of GABAergic tonic currents in thalamic relay neurons is sufficient to produce SWDs in wild-type rats, and multiple rodent models of absence epilepsy (GAERS, stargazer, lethargic, tottering) express *increases* in thalamic inhibitory tonic currents (Fariello & Golden, 1987; Crunelli et al., 2011; Errington et al., 2011). The present study, however, demonstrates that this link must be expanded to include tonic inhibition in cortical neurons. Using continuous video-EEG monitoring and selective pharmacological manipulation of cortical tonic inhibition, we show that *decreasing* cortical inhibitory tonic currents is also ‘sufficient’ to produce SWDs in wild-type (RR) mice, and that *rescuing* the lost cortical tonic currents in RQ mice suppresses SWD expression.

## RESULTS

### RQ mice express SWDs and absence epilepsy

The γ2R43Q mutation confers absence seizures and generalized EEG SWDs in humans^3^ and knock-in (RQ) mice (Tan et al, 2007). Figure 1 illustrates bilateral, synchronous (∼6 Hz) SWDs in a RQ mice using continuous EEG and electromyogram (EMG) recordings. Quantification was done off-line after recordings were completed. A SWD ‘bout’ was classified as two or more individual SWD events occurring <30 seconds apart. SWDs were assessed for individual event duration, inter-bout-intervals (IBI), events per bout, and bout duration. EEG and EMG recordings from one RQ mouse during a SWD bout are presented (Fig. 1A & 1B), along with quantified SWD assessment for three different RQ mice (Fig. 1C). All RQ mice assessed with EEG and EMG monitoring presented synchronized SWDs across all EEG leads with coincident cessation of EMG activity. No SWDs were observed in naïve wild-type (RR) mice. All values for all groups and conditions and measures are presented in Table 1.

**Table 1.**
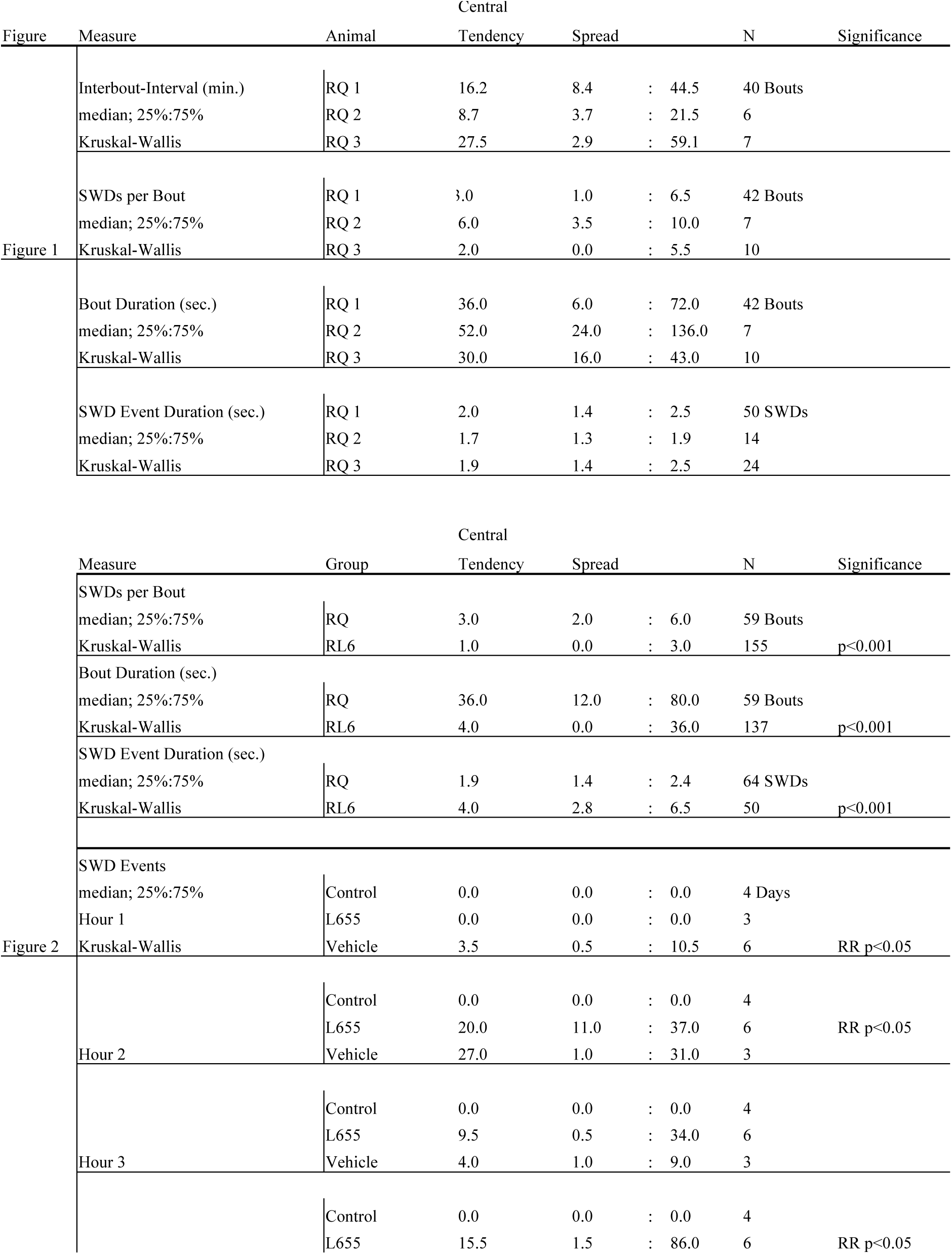
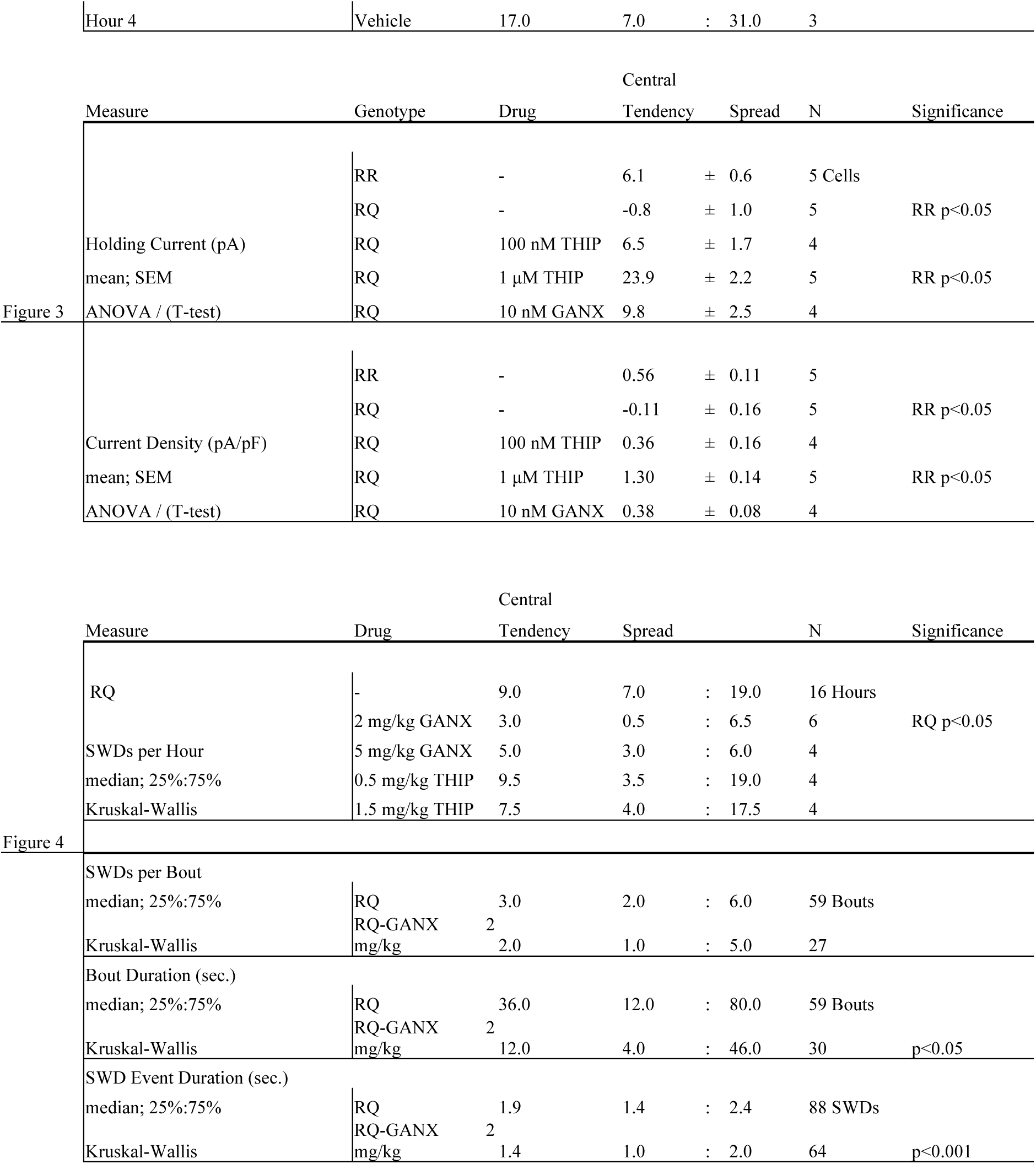
– Measures and statistics. All measures (mean or median), spreads (SEM or 25%:75%), N (number of samples) and statistical significance (ANOVA or Kruskal-Wallis) for all groups and conditions compared are presented according to associated Figure.

**Figure 1.**
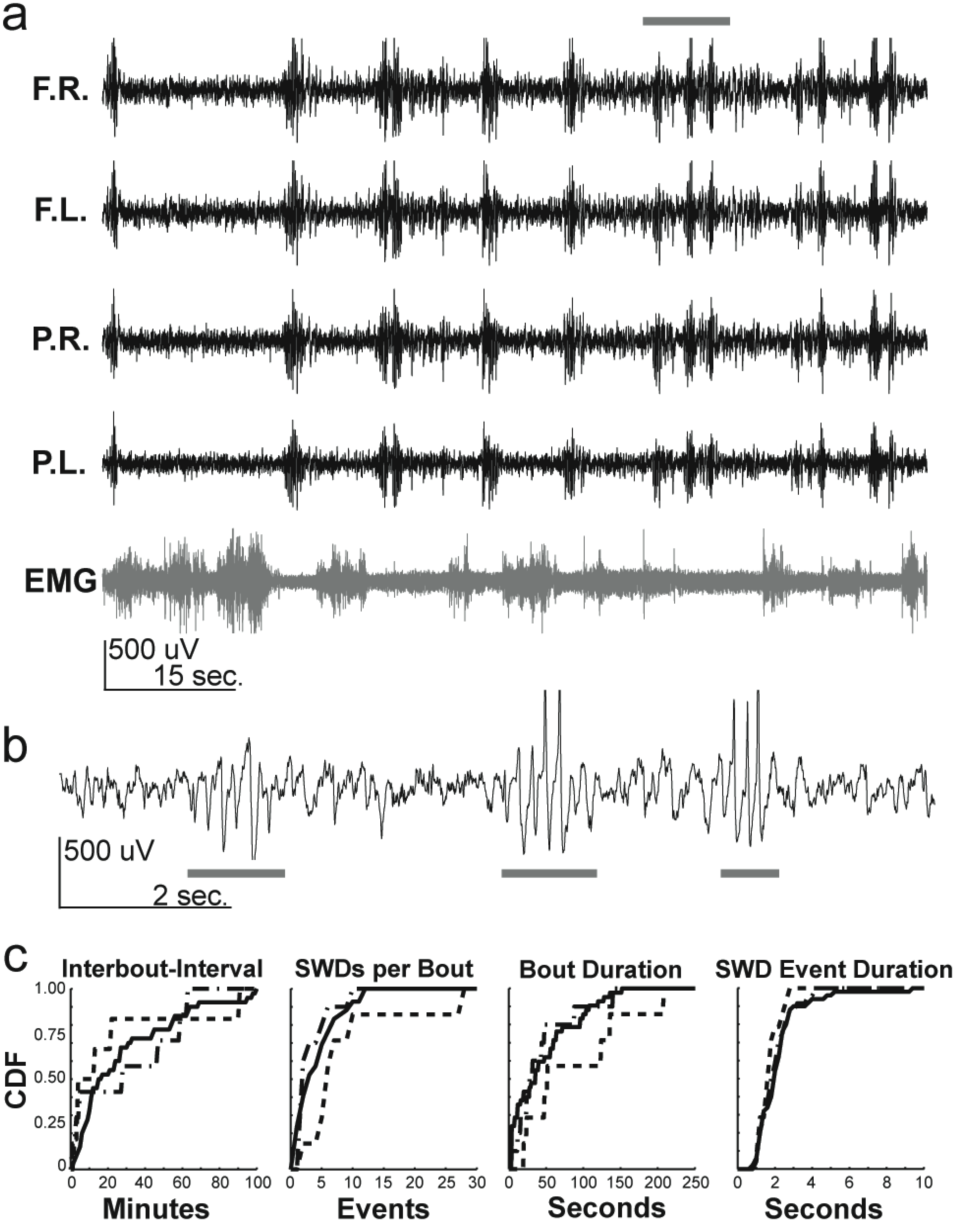
RQ mice express SWDs associated with absence epilepsy. A) Electroencephalogram (EEG) recording of an RQ mouse. Top trace to bottom trace: frontal right cortex (F.R.); frontal left cortex (F.L.); parietal right cortex (P.R.); parietal left cortex (P.L.); electromyogram (EMG). Note the brief yet frequent (∼11 times during the 1.5 minute trace) synchronized events that occur across all EEG leads during the absence of signal in the EMG. B) Expanded F.R. EEG recording from grey bar in A (10 seconds). Note the brief ∼6 Hz SWD events (grey bars) that occur 3 times during the 10-second trace. C) Cumulative distributions from three different RQ mice (solid, dashed, and dash-dotted lines represent each mouse) show similar characteristics from all animals for interbout-interval, SWDs per bout, bout duration and SWD event duration. SWDs were not observed in litter-mate control mice (not shown).

### Blocking cortical tonic inhibition produces SWDs in wild-type mice

It has been demonstrated that a positive correlation between SWDs and thalamic inhibitory tonic currents exists (Cope et al., 2009), leading to the conclusion that enhanced GABAergic tonic inhibition is “necessary and sufficient” condition to cause typical absence epilepsy (Crunelli et al., 2011; Errington et al., 2011). In sharp contrast, here we demonstrate that altering thalamic tonic inhibition is not necessary for SWD generation, and furthermore, blockade of cortical tonic inhibition is sufficient to produce SWDs (Fig. 2). Previously we reported that inhibitory tonic currents in somatosensory cortical layer II/III principal neurons are generated by α5 subunit-containing GABA_A_ receptors and that this current is blocked by the α5 subunit-selective inverse agonist L655,708 (L655) (Mangan et al., 2013). Intraperitoneal (i.p.) administration of L655, at a concentration (2 mg/kg) known to bind the majority of α5 subunit-containing receptors (Atack et al., 2006), produced SWDs (∼6 Hz) in RR mice (RRL6) that are electrographically similar to SWDs seen in RQ mice (Fig. 2A & 2B). However, L655 induced SWDs (L6-SWDs) display fewer events per bout (p<0.001) and shortened bout durations (p<0.001) compared to RQ, while individual L6-SWD event duration was longer (p<0.001). Also noteworthy is the appearance of SWDs 3 days after the last L655 injection (Fig. 2D, hour 1 vehicle), suggesting lingering plasticity following initial insult and SWD induction.

**Figure 2.**
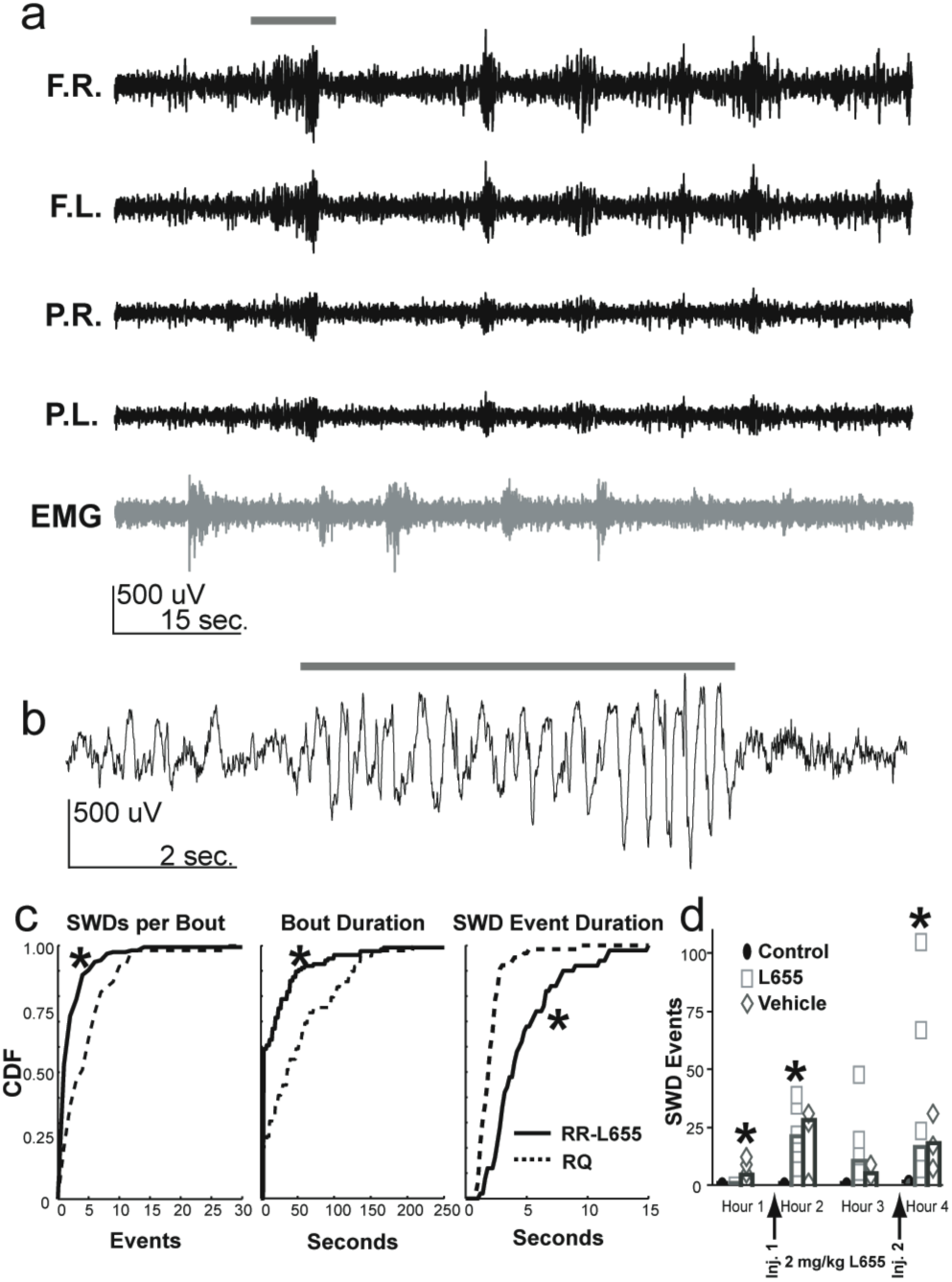
Blocking cortical tonic inhibition produces SWDs in wild-type mice. A) Electroencephalogram (EEG) recording of a wild-type (RR) mouse i.p. injected with 2 mg/kg of the GABA_A_ receptor α5 subunit-selective inverse agonist L655,708 (RR-L655). Similar to RQ mice, note the brief yet frequent (∼6 times during the 1.5 minute trace) synchronized events that occur across all EEG leads during the absence of signal in the EMG. B) Expanded F.R. EEG recording from grey bar in A (10 seconds) displays prolonged ∼6 Hz SWD event (grey bar). C) Cumulative distributions show RR-L655 mice display significantly less SWDs per bout (p<0.05), shorter bout durations (p<0.05), yet longer SWD event durations (p<0.05) than RQ mice. D) Quantification of SWD events shows that RR-L655 mice did not display SWDs prior to L655,708 injection (Hour 1), but did show SWDs after each hour of injection (Hour 2, p<0.05; Hour 4, p<0.05). Interestingly, SWDs were still present in RR-L655 mice 3 days after the last L655,708 treatment (vehicle: Hour 1, p<0.05).

### GABA_*A*_ receptor δ subunit-selective agonists rescue tonic inhibition in RQ cortical principal neurons

Although RQ principal cortical neurons lack inhibitory tonic currents (Mangan et al., 2013), these neurons also display an inhibitory conductance in response to selective δsubunit-associated GABA_A_ receptor agonists (1 μM THIP (Cope et al., 2005; Adkins et al., 2001; Mangan et al., 2013), 30 nM allopregnanolone (ALLO) (Fodor et al., 2005; Rajasekaran et al., 2010; Mangan et al., 2013). This finding is consistent with the presence of latent δ subunit-containing GABA_A_ receptors in RQ cortical neurons and suggests that the lost tonic inhibition in these neurons can be rescued. We used whole-cell patch-clamp recordings to titrate a concentration of selective δ subunit-containing GABA_A_ receptor agonists that rescued wild-type tonic inhibition levels in RQ cortical neurons. We found that a low concentration (10 nM) of Ganaxolone (GANX) (Fig. 3B), a synthetic neuroactive steroid related to ALLO (Citraro et al., 2006), activates a latent inhibitory conductance in RQ cortical neurons equal to the inhibitory tonic current observed in RR cortical neurons.

**Figure 3.**
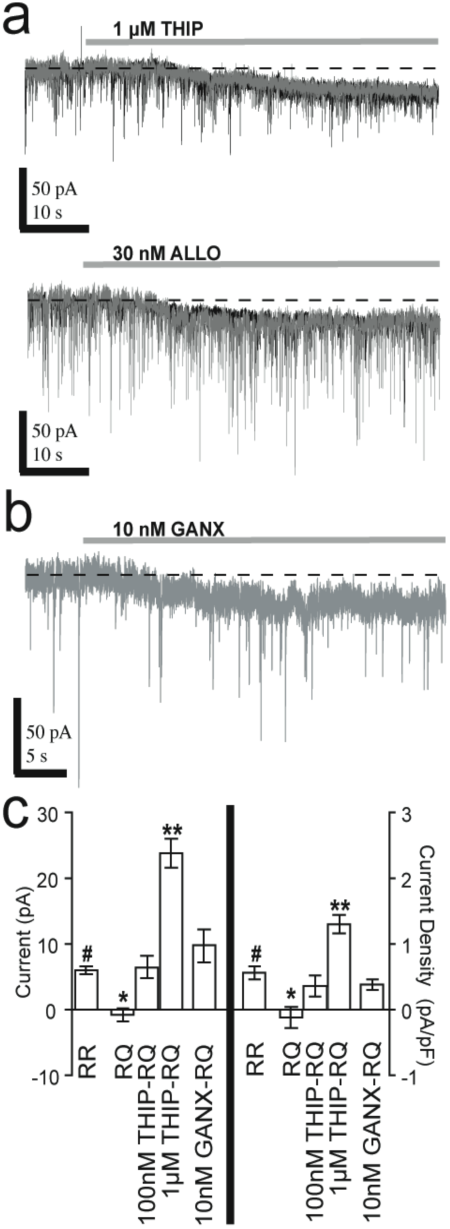
GABA_A_ receptor δ subunit-selective agonists rescue tonic inhibition in RQ cortical neurons. A) Example voltage-clamp traces for RR (black-behind) and RQ (grey-front) cortical layer II/III cell recordings during 1 μM THIP (top) and 30 nM allopregnanolone (ALLO: bottom) treatments. Both GABA_A_ receptor μ subunit-selective agonist treatments induce indistinguishable current amplitudes and densities in RQ compared to RR. B) Example voltage-clamp trace for RQ cortical layer II/III cell recording during a 10 nM ganaxolone (GANX) treatment also shows an increase in the holding current, similar to THIP and ALLO. C) Tonic current amplitude (left y-axis) and density (right y-axis) quantifications reveal normal RR tonic inhibition levels can be rescued in RQ with 100 nM THIP and 10 nM GANX treatments, whereas 1 μM THIP treatment in RQ produces 2-4 times more holding current amplitude (p<0.05) and density (p<0.05) than that observed in untreated RR neurons.

### Rescuing cortical tonic inhibition attenuates SWDs in RQ mice

Low doses of selective δ subunit-containing GABA_A_ receptor agonists rescue tonic inhibition in RQ principal cortical neurons (Fig. 3B). Using video-EEG monitoring, we investigated if treatment with these δ subunit-selective GABA_A_ receptor agonists (GANX or THIP) could ameliorate the SWDs observed in RQ mice.

RQ mice were i.p. injected twice a day with GANX or THIP for 4 out of 7 days (for drug schedule see Fig. 4A). Multiple concentrations of GANX (2 and 5 mg/kg) and THIP (0.5 and 1.5 mg/kg) were tested for their ability to suppress SWD expression. Only the lowest concentration (2 mg/kg) of GANX was significantly (p<0.05) effective in decreasing RQ-SWD expression (Fig. 4B). This low dose of GANX treatment also decreased bout duration (p<0.05) and event duration (p<0.001), but did not affect the number of SWDs per bout. The selective efficacy of the low GANX dose (2 mg/kg), which is half the ED_50_ dose that protects against partial seizures (Gasior et al., 1999; Kaminski et al., 2004), suggests that the mechanism that diminishes SWDs in RQ mice involves activation of latent δ subunit-containing GABA_A_ receptors in cortical neurons.

**Figure 4.**
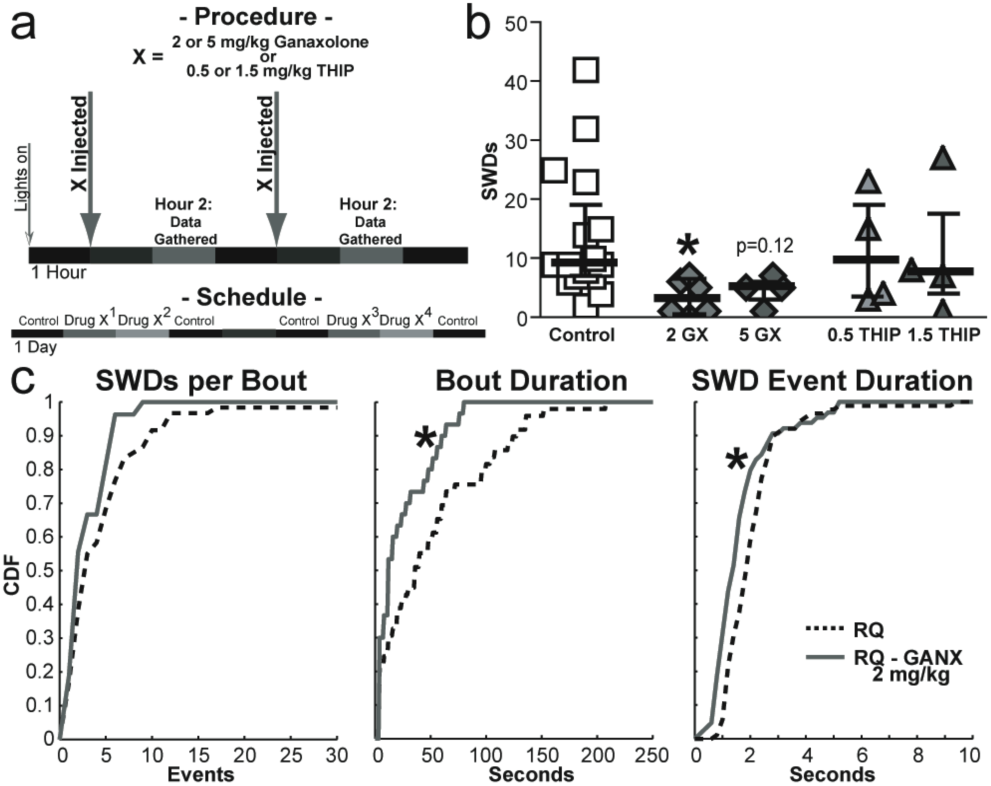
Rescuing cortical tonic inhibition alleviates SWDs in RQ mice. A) Schematic depicting administration times and drug schedule investigating 4 drug-treatment conditions in RQ mice. GANX (2 and 5 mg/kg) or THIP (0.5 and 1.5 mg/kg) solutions were i.p. injected in RQ mice twice a day for 4 out of 7 days. B) RQ SWD event quantification during the second hour after drug administration shows the 2 mg/kg GANX treatment decreased SWD expression compared to control hours (p<0.05). C) Cumulative distributions of RQ SWD activity after 2 mg/kg GANX treatment shows that bout (p<0.05) and SWD event (p<0.05) durations are decreased after treatment.

## DISCUSSION

The major findings from this study are that the loss (RQ) (Fig. 1) or decrease (RR-L655) (Fig. 2) of cortical tonic inhibition results in a SWD-expressing phenotype, and the pharmacological replacement of cortical tonic inhibition (RQ-GANX: Fig. 3C) suppresses SWD expression (Fig. 4). These findings are consistent with the conclusion that the amount of cortical tonic inhibition regulates SWD expression. Thus, considering previous findings along with our research suggests the causal link between absence epilepsy and inhibitory tonic currents is at least bidirectional: *increased* thalamic tonic inhibition (Cope et al., 2009) or *decreased* cortical tonic inhibition; both can lead to epileptiform activity.

The link between absence seizures and increased δ subunit-associated GABA_A_ receptor activation in thalamic relay neurons is well established (Fariello & Golden, 1987; Crunelli et al., 2011; Errington et al., 2011). The current hypothesis from this evidence is that persistent hyperpolarization of thalamic relay neurons (Cope et al., 2005, 2009) makes these neurons more susceptible to rhythmic bursting and insensitive to sensory input, which is considered to be a necessary condition for SWD generation (Crunelli et al., 2011; Errington et al., 2011). Consistent with this hypothesis, ethosuximide and valproic acid, two different T-type Ca^2+^ channel blockers, decrease thalamic relay bursting and are currently the main treatment options for absence epilepsy. However, the efficacy of either drug for this condition is at only ∼55% (Glauser et al., 2010). The evidence presented in this study suggests a second classification for absence seizure generation that is separate from altered thalamic activity, and could apply to at least a portion of the remaining ∼45% of patients that are currently not treatable by the main treatment options.

*In vitro* examination of thalamic relay bursting behaviors in thalamocortical mouse brain slices detected a decrease or no change in bursting behaviors compared to control for RQ and L655-treated (RR) brain slices, respectively (Mangan et al., 2013). Additionally, RQ thalamic relay neurons display no inhibitory tonic currents (Mangan et al., 2013). These results suggest that neither increased thalamic tonic inhibition nor the resulting increased susceptibility to rhythmic bursting is essential for SWD generation. Furthermore, the SWDs expressed endogenously in WAG/Rij rats is quenched by local application of ganaxolone or allopregnanolone into somatosensory cortex, a treatment that would not directly affect thalamic relay neuron tonic inhibitory levels or bursting susceptibility (Citraro et al., 2006). It is likely, however, that this treatment to WAG/Rij rats increased cortical tonic inhibitory levels, further suggesting that cortical inhibitory levels regulate SWD expression.

Our findings suggest that SWDs are linked to general cortical tonic inhibition levels and not to a specific tonic current-associated GABA_A_ receptor subtype (α5 or δ). Rescuing RQ cortical tonic inhibition via activation of δ-subunit-associated GABA_A_ receptors with GANX, and the subsequent decrease in SWD expression (Fig 4), indicates that SWD expression can be regulated by δ-subunit-associated tonic inhibition. Conversely, the selective decrease / block of α5 subunit-associated inhibition (RR-L655), which results in SWD expression (Fig. 2), indicates that SWD expression can also be regulated by α5 subunit-associated tonic inhibition.

We found evidence of long-lasting aberrant thalamocortical function after inducing SWDs with L655 in wild-type mice. Mice that were injected twice a day for 2 consecutive days with L655 still displayed SWDs 3 days after the last injection (Fig. 2D: vehicle, Hour 1, p<0.05). Given that the half-life of L655 in multiple animal models (rat, dog, rhesus monkey) is 0.3 – 0.5 hours (Atack et al., 2009), this result suggests lingering pro-epileptic plasticity of the thalamocortical circuit follows initial seizure insult. Similar pro-seizure susceptibility has been noted in other epilepsy-induced animal models (Yu et al., 2013; Peng et al., 2004; Houser & Esclapez, 2003). Getting the earliest possible therapeutic intervention should be a high priority for individuals suffering from absence epilepsy.

### Optimal Tonic Inhibition

GANX is the only neurosteroid evaluated as an anti-epileptic drug in humans (Monaghan et al., 1999; Nohria et al., 2010; Reddy and Rogawski 2012). It has been clinically studied for the treatment of infantile spasms (Kerrigan et al., 2000) and shown to be effective, with minimal side effects (sedation), as a treatment for catamenial epilepsy (Reddy and Rogawski, 2009) and partial seizures (Laxer et al., 2000) in adults. However, investigation of GANX in animal models of absence epilepsy (PTZ, GHB) uncovered that it exacerbates absence seizures and can even produce SWDs in wild – type rats when administered at ≥ 20 mg/kg (Snead III; 1998). Thus, how can we suggest GANX can be effective as a treatment for absence epilepsy?

Neurosteroids activate GABA_A_ receptors directly but are known to produce the largest magnitude effects at δ subunit-containing GABA_A_ receptors and are selective for this receptor subtype only at lower concentrations (Reddy and Rogawski, 2009). Our current results suggest a dichotomy of effects for neurosteroids in the CNS: higher concentrations result in general sedation and SWD generation or exacerbation, and lower levels produce normal functionality and ameliorate SWDs in RQ mice. Similarly, we suggest a concentration dependent consideration of thalamocortical tonic inhibition in regards to SWDs and absence seizure generation.

Evaluation of polygenic (GAERS, stargazer, lethargic) and pharmaco-induced (GHB, PTZ) rodent models of absence epilepsy provide substantial evidence that too much thalamic tonic inhibition triggers SWDs (Cope et al., 2009; Snead III, 1998). Equally, here we present a novel absence animal model (L655) and treatment for a known absence animal model (RQ) that indicate reduced cortical tonic inhibition results in SWDs. These findings suggest that an optimum level of tonic inhibition in the thalamocortical circuit is required for normal functioning and that deviation from this optimum in either direction results in aberrant thalamocortical function, SWDs and absence seizures.

## ACKNOWLEDGEMENTS

This work was supported by grants from the Epilepsy Foundation (K.P.M., M.V.J.) and NIH (NS046378, NS075366 to M.V.J.).

## AUTHOR CONTRIBUTIONS

K.P.M. and M.V.J. jointly conceived and designed all experiments for the study with guidance from S.P. and C.C.; K.P.M. and A.B.N. performed experiments; K.P.M. and M.V.J. analyzed data and wrote the manuscript.

## METHODS

### EEG implantation and monitoring of SWDs

The present study used male and female wild-type Harlan C57BL/6J-OlaHsd and ?2R43Q knock-in mice bred into a background of Harlan C57BL / 6J-OlaHsd mice. Behavioral and electrographic markers of absence epilepsy in these animals were confirmed by video-EEG monitoring. Surgery and electrode implantation are described in Nelson et al. (2013). Briefly, P24 mice were implanted, under isoflurane anesthesia (1% – 2% in 100% O_2_), for chronic EEG recordings with gold plated miniature screw electrodes over the right and left frontal and parietal cortices, and one over the cerebellum as reference. Two vinyl-coated braided stainless steel wire electrodes were placed in the nuchal muscle for electromyogram (EMG) recording of muscle activity. All electrodes were gathered into a flexible cable and connected to the Multichannel Neurophysiology Recording system (Tucker-Davis Technologies, TDT, Alachua, FL, USA). EEG and EMG signals were collected continuously at a sampling rate of 256 Hz (digitally filtered between 0.1 and 100 Hz). Continuous EEG recordings with occasional video monitoring were made and SWDs were scored off-line. Animals were given a 3-day recover period after surgery before SWD scoring began. A SWD event was defined as a brief (∼2 seconds long) ∼6 Hz signal synchronized across all EEG leads, with a corresponding lack of signal in the EMG lead. Only SWD events that occurred >2 min from slow-wave-sleep periods were used for quantification. SWD event durations were measured from the first synchronized positive peak signal to the last synchronized positive peak within an event. SWD “bouts” were defined as groups of SWD events separated from other events by <30 seconds. Interbout-intervals were defined as the time between the beginnings of consecutive bouts.

All animal procedures followed the National Institutes of Health Guide for the Care and Use of Laboratory Animals and facilities were reviewed and approved by the IACUC of the University of Wisconsin-Madison, and were inspected and accredited by AAALAC.

### Drugs and Injection Schedule

L655,708 (L9787), GANX (G7795) and THIP (T101) were all obtained from Sigma (St. Louis, MO). L655 and GANX were dissolved in a 30% DMSO-saline solution (v/v), while THIP was dissolved in 100% saline. Mice were intraperitoneally (i.p.) injected with 2 mg/kg doses of L655, 2 and 5 mg/kg doses of GANX, or 0.5 and 1.5 mg/kg doses of THIP. 160 uL of solution was injected for each drug. L655 was administered to RR mice 2 and 4 hours after lights out (Fig 2) for 2 consecutive days beginning 5 days after surgery. These mice were not injected for the subsequent 2 days, but were given vehicle injections on day 9. Ganxolone or THIP injections were administered to RQ mice 1 and 4 hours after lights out (Fig. 4). Drug injections for RQ mice began on day 5 after surgery and consisted of 2 injections of one drug and dose, with a different drug and dose for days 6, 10, and 11. No injections were given to RQ mice on days 7-9.

### Whole-cell Patch Clamp Experiments

Horizontal slices (400 μm) were prepared from the brains of RR and RQ mice of either sex (16 – 26 days old). All procedures were approved by the University of Wisconsin Institutional Animal Care and Use Committee. Mice were anesthetized with isoflurane, decapitated, and the brain was removed and placed in ice-cold cutting solution containing (in mM): 125 NaCl, 25 NaHCO_3_, 2.5 KCl, 1.25 NaH_2_PO_4_, 0.5 CaCl_2_, 3.35 MgCl_2_, 25 D-Glucose,13.87 M sucrose, and bubbled with 95% O_2_ and 5% CO_2_. Slices were cut using a vibratome (Leica VT 1000S, Global Medical Imaging; Ramsey, MN) and placed in an incubation chamber containing standard artificial cerebrospinal fluid (aCSF) (in mM): 125 NaCl, 25 NaHCO_3_, 2.5 KCl, 1.25 NaH_2_PO_4_, 2 CaCl_2_, 1 MgCl_2_, 25 D-Glucose, at room temperature for 1 hour before being used for recordings. Whole cell patch-clamp recordings were made from somatosensory cortical layer II/III pyramidal cells, visualized using an upright differential interference contrast microscope (Axioskop FS2, Zeiss; Oberkochen, Germany). Patch pipettes were pulled from thin-walled borosilicate glass (World Precision Instruments; Sarasota, FL) with a resistance of 3-5 MΩ when filled with intracellular solution containing (in mM): 140 KCl, 10 EGTA, 10 HEPES, 20 phosphocreatine, 2 Mg_2_ATP, 0.3 NaGTP (pH 7.3, 310 mOsm). Voltage-clamp (-60 mV) recordings were made in a submerged chamber at room temperature using a MultiClamp 700B amplifier (Axon Instruments; Foster City, CA), filtered at 4 kHz and digitized at 10 kHz using a Digidata 1322A analog-digital interface (Axon Instruments). Data were acquired to a Macintosh G4 (Apple Computer; Cupertino, CA) using Axograph X v1.1.4 (Molecular Devices; Sunnyvale, CA).

Data segments (30 s) just prior to and 90 s after drug administration were analyzed to quantify inhibitory tonic currents. All-point amplitude histograms were computed for each segment, and fit with a Gaussian function only to the outward current portions relative to the peak in order to omit components arising from inward phasic mIPSCs^39^. Tonic current was calculated as the difference between the fitted Gaussian means before and after (100 or 1 μM) THIP, (30 nM) ALLO, or (10 nM) GANX administration. Current density (pA/pF) was calculated by dividing the current by cell capacitance. Bicuculline (100 μM) was added at the conclusion of at least one experiment for each drug tested to verify full current block and, thus, only GABAergic contribution.

### Statistics

The Kruskal-Wallis test of medians was used to compare multiple groups with a Dunn’s post-hoc evaluation. Tonic current amplitude and density data were normally distributed, thus an ANOVA was used to compare multiple groups with a Bonferroni post-hoc evaluation. Matlab and Prism software was used.

## REFERENCES

Adkins, C.E., et al. alpha4beta3delta GABA(A) receptors characterized by fluorescence resonance energy transfer-derived measurements of membrane potential. J Biol Chem 276, 38934–38939 (2001).

Atack J.R. et al. L-655,708 enhances cognition in rats but is not proconvulsant at a dose selective for alpha5-containing GABAA receptors. Neuropharmacology 51(6):1023–9 (2006).

Atack J.R. et al. The plasma – occupancy relationship of the novel GABAA receptor benzodiazepine site ligand, a5IA, is similar in rats and primates. BR J Pharmaco 157: 796–803 (2009).

Bazyan A.S., van Luijtelaar G. Neurochemical and behavioral features in genetic absence epilepsy and in acutely induced absence seizures. ISRN Neurol May 7 (2013).

Braudeau J. et al. Chronic treatment with a promnesiant GABAA α5-selective inverse agonist increases immediate early genes expression during memory processing in mice and rectifies their expression levels in a down syndrome mouse model. Adv Pharmacol Sci 153218 (2011).

Chen S.D., Yeh K.H., Huang Y.H., Shaw F.Z. Effect of intracranial administration of ethosuximide in rats with spontaneous or pentylenetetrazol-induced spike-wave discharges. Epilepsia 52(7): 1311–8 (2011).

Citraro R., et al. Effects of some neurosteroids injected into some brain areas of WAG/Rij rats, an animal moel of generalized absence epilepsy. Neuropharm 50: 1059–71 (2006).

Cope D.W., Hughes S.W., Crunelli V. GABAA receptor-mediated tonic inhibition in thalamic neurons. J Neurosci 25, 11553–11563 (2005).

Cope, D.W., et al. Enhanced tonic GABAA inhibition in typical absence epilepsy. Nat Med 15, 1392–1398 (2009).

Crunelli V., Cope, D.W., Terry J.R. Transition to absence seizures and the role of GABA(A) receptors. Epilepsy Res 97, 283–289 (2011).

Errington A.C., Cope D.W., Crunelli V. Augmentation of Tonic GABA(A) Inhibition in Absence Epilepsy: Therapeutic Value of Inverse Agonists at Extrasynaptic GABA(A) Receptors. Advances in pharmacological sciences 2011, 790590 (2011).

Eugene E., et al. GABA(A) receptor gamma 2 subunit mutations linked to human epileptic syndromes differentially affect phasic and tonic inhibition. J Neurosci 27, 14108–14116 (2007).

Fariello R.G., Golden G.T. The THIP-induced model of bilateral synchronous spike and wave in rodents. Neuropharmacology 26, 161–165 (1987).

Fedi, M., et al. Intracortical hyperexcitability in humans with a GABAA receptor mutation. Cereb Cortex 18, 664–669 (2008).

Fodor L., Biro T., Maksay G. Nanomolar allopregnanolone potentiates rat cerebellar GABAA receptors. Neurosci Lett 383, 127–130 (2005).

Gasior M et al. Acute and chronic effects of the synthetic neuroactive steroid, ganaxolone, against the convulsive and lethal effects of pentylenetetrazol in seizure-kindled mice: comparison with diazepam and valproate. Neuropharmacology 39(7): 1184–96 (2000).

Glauser TA et al. (2010) Childhood Absence Epilepsy Study Group. Ethosuximide, valproic acid, and lamotrigine in childhood absence epilepsy. N Engl J Med 362(9):790–9.

Hales, T.G., et al. The epilepsy mutation, gamma2(R43Q) disrupts a highly conserved inter-subunit contact site, perturbing the biogenesis of GABAA receptors. Mol Cell Neurosci 29, 120–127 (2005).

Herd M.B. et al. Balfour D.J., Lambert J.J., Belelli D. Inhibition of thalamic excitability by 4,5,6,7-tetrahydroisoxazolo[4,5-c]pyridine-3-ol: a selective role for delta-GABA(A) receptors. Eur J Neurosci 29(6):1177–87 (2009).

Houser C.R., Esclapez M Downregulation of the alpha5 subunit of GABAA receptor in the pilocarpine model fo temporal lobe epilepsy. Hippocampus 13(5): 633–45.

Kaminski R.M., Livingood M.R., Rogawski M.A. Allopregnanolone analogs that positively modulate GABAA receptors protect against partial seizures induced by 6-Hz electrical stimulation in mice. Epilepsia 45(7): 864–7 (2004).

Kang, J.Q., Macdonald, R.L. The GABAA receptor gamma2 subunit R43Q mutation linked to childhood absence epilepsy and febrile seizures causes retention of alpha1beta2gamma2S receptors in the endoplasmic reticulum. J Neurosci 24, 8672–8677 (2004).

Kerrigan J.F. et al. Ganaxolone for treating intractable infantile spasms: a multicenter, open-label, add-on trial. Epilepsy Res 42: 133 – 139 (2000).

Kleschevnikov A.M. et al. Mobley W.C. Deficits in cognition and synaptic plasticity in a mouse model of Down syndrome ameliorated by GABAB receptor antagonists. J Neurosci 32(27): 9217–27 (2012).

Laxer K. et al. Assessment of ganaxolone’s anticonvulsant activity using a randomized, double-blind, presurgical trial design. Ganaxolone Presurgical Study Group. Epilepsia 41: 1187 – 1194 (2000).

Littjohann A., Zhang S., de Peijper R., van Luijtelaar G. Electrical stimulation of the epileptic focus in absence epileptic WAG/Rij rats: assessment of local and network excitability. Neuroscince 188:125–34 (2011).

Littjohann A., van Luijtelaar G. The dynamics of corticothalamo-cortical interactions at the transition from pre-ictal to ictal LFPs in absence epilepsy. Neurobiol Dis 47(1):49–60 (2012).

Mangan, K.P. (2013) GABAergic Tonic Inhibition, Thalamocortical Function, and Absence Epilepsy. Ph.D. Thesis, University of Wisconsin – Madison

Manning J.P., Richards D.A., Leresche N., Crunelli V., Bowery N.G. Cortical-area specific block of genetically deteremined absence seizures by ethosuximide. Neuroscience 123(1): 5–9. (2004).

Martínez-Cué C. et al. Reducing GABAA a5 receptor-mediated inhibition rescues functional and neuromorphological deficits in a mouse model of down syndrome. J Neurosci 33(9): 3953–66 (2013).

McCormick, D.A., Contreras, D. On the cellular and network bases of epileptic seizures. Annu Rev Physiol 63, 815–846 (2001).

Meeren H.K., Pijn J.P., Van Luijtelaar E.L., Coenen A.M., Lopes da Silva F.H. Cortical focus drives widespread corticothalamic networks during spontaneous absence seizures in rats. J Neurosci 22, 1480–1495 (2002).

Meeren H., van Luijtelaar G., Lopes da Silva F., Coenen A. Evolving concepts on the pathophysiology of absence seizures: the cortical focus theory. Archives of neurology 62, 371–376 (2005).

Monaghan E.P., McAuley J.W., Data J.L. Ganaxolone: a novel positive allosteric modulator of the GABAA receptor complex for the treatment of epilepsy. Expert Opin Investig Drugs 8:1663 – 1671 (1999).

Nelson A.B., Faraguna U., Zoltan J.T., Tononi G., Cirelli C. Sleep patterns and homeostatic mechanisms in adolescent mice. Brain Sci 3, 318–343 (2013).

Nohria V. et al. Ganaxolone. In: Bialer M, et al., editors. Epilepsy Res; Progress report on new antiepileptic drugs: a summary of the Tenth Eilat Conference (EILAT X); p. 89–124 (2010).

Panayiotopoulos, C.P. Typical absence seizures and related epileptic syndromes: assessment of current state and directions for future research. Epilepsia 49, 2131–2139 (2008).

Peng Z., Huang C.S., Stell B.M., Mody I., Houser C.R. Altered expression of the delta subunit of the GABAA receptor in a mouse model of temporal labe epilepsy. J Neurosci 24(39): 8629–39.

Pinault D. Cellular interactions in the rat somatosensory thalamocortical system during normal epileptic 5 – 9Hz oscillations. J Physio 552(3): 881–905 (2003).

Polack P.O., et al. Deep layer somatosensory cortical neurons initiate spike-and-wavedischarges in a genetic model of absence seizures. J Neurosci 27(24): 6590–9. (2007).

Rajasekaran, K., Joshi, S., Sun, C., Mtchedlishvilli, Z. & Kapur, J. Receptors with low affinity for neurosteroids and GABA contribute to tonic inhibition of granule cells in epileptic animals. Neurobiology of disease 40, 490–501 (2010).

Reddy D.S., Rogawski M.A. Neurosteroids replacement therapy for catamenial epilepsy. Neurotherapeutics 6(2): 392–401 (2009).

Reddy D.S., Rogawski M.A. Neurosteroids – endogenous regulators of seizure susceptibility and role in the treatment of epilepsy. In Jasper’s Basic Mechanisms of the Epilepsies; 4th edition. Bethesda (MD) (2012).

Sancar, F. & Czajkowski, C. A GABAA receptor mutation linked to human epilepsy (gamma2R43Q) impairs cell surface expression of alphabetagamma receptors. J Biol Chem 279:47034–47039 (2004).

Sitnikova E., van Luijtelaar G. Cortical control of generalized absence seizures: effect of lidocaine applied to the somatosensory cortex in WAG/Rij rats. Brain Res 1012(1-2): 127–37 (2004).

Snead III O.C. Ganaxolone, a selective, high-affinity steroid modulator of the gamma-aminobutyric acid-A receptor, exacerbates seizures in animal models of absence. Annals of Neurology 44(4): 688 – 691 (1998).

Steriade M., Contreras D. Spike-wave complexes and fast components of cortically generated seizures. I. Role of neocortex and thalamus. J Neurophysiol 80(3): 1439–55 (1998).

Tan, H.O., et al. Reduced cortical inhibition in a mouse model of familial childhood absence epilepsy. Proc Natl Acad Sci U S A 104: 17536–17541 (2007).

van Luijtelaar G., Sitnikova E. Global and focal aspects of absence epilepsy: the contribution of genetic models. Neurosci Biobehav Rev 30(7): 983–1003 (2006).

van Luijtelaar G., Sitnikova E., Littjohann A. On the origin and suddenness of absences in genetic absence models. Clin EEG Neurosci 42(2): 83–97 (2011).

Wallace, R.H., et al. Mutant GABA(A) receptor gamma2-subunit in childhood absence epilepsy and febrile seizures. Nat Genet 28: 49–52. (2001).

Yu T., et al. Surgical treatment of hyermotor seizures originating from the temporal lobe. Seizure S1059–1311 (2013).

